# An abundant myeloid progenitor shapes neonatal hematopoiesis of naked mole-rats

**DOI:** 10.1101/2021.08.01.454658

**Authors:** Stephan Emmrich, Alexandre Trapp, Andrei Seluanov, Vera Gorbunova

## Abstract

The naked mole-rat became an attractive model for biomedical research due to its exceptional longevity and resistance to disease, yet the developmental properties of naked mole-rat stem and progenitor compartments are poorly understood. Here we report a high frequency bone marrow progenitor, which we termed TC90, exclusively present in neonatal animals. TC90s were marked by an immunophenotype resembling adult naked mole-rat common myeloid progenitors (CMP) with mature neutrophilic Thy1.1-antigen levels and showed a clear differentiation bias and transcriptome signature towards myelopoiesis. TC90s overexpressed both CXCL12 and its alternative receptor CXCR7, thereby abrogating their homing capacity and produced multiple secreted matrix components related to neonatal bone marrow niche formation. We hypothesize that the TC90 cell population protects newborns during the critical transition at birth and shortly beyond by boosting a supply of innate immune cells. In summary, our results identify a transient cell stage during postnatal hematopoiesis unique to naked mole-rats.

## Introduction

The naked mole-rat (*Heterocephalus glaber*) has become an attractive animal model in aging research due to its longevity and resistance to disease. Naked mole-rats are subterranean eusocial rodents found in grasslands of East Africa. This remarkable rodent with the size of a mouse has a maximum lifespan of over 39 years in captivity and does not display increased mortality during aging^1, 2^. Previously we discovered that high molecular weight hyaluronic acid is responsible for the resistance of naked mole-rats to solid tumors^3^, while further unique molecular adaptations such as a split 28S rRNA and a fusion isoform in the INK4A/ARF locus confer improved cellular responses towards senescence and transformation^4, 5^. We created a naked mole-rat hematlas by employing cross-reactive antibodies for fluorescence-activated cell sorting (FACS), purifying hematopoietic stem and progenitor cell (HSPC) subsets and using colony formation, xenotransplantation and Cellular Indexing of Transcriptomes and Epitopes by Sequencing (CITE-Seq) to link single-cell transcriptomic cell type annotation with FACS immunophenotypes^6^. Hallmarks of naked mole-rat hematopoiesis are high basal levels of neutrophil granulocytes (GC) accompanied by reduced B-lymphopoiesis, splenic erythropoiesis as major route of red blood cell production, prolonged cell cycle duration of hematopoietic cells and a primitive HSPC compartment marked by CD34, THY1 and TM4SF1. Moreover, we revealed the presence of an ectopic cervical thymus along with a delay of immunosenescence and thymic involution in naked mole-rats, and showed that naked mole-rat T-lymphopoiesis results in strongly reduced CD8^+^ cytotoxic T cell frequencies^7^.

Every second, an adult human produces nearly 2 million red blood cells^8^. This enormous turnover rate, required to sustain life in all vertebrates, is accomplished unilaterally by hematopoietic stem cells (HSCs). HSCs are self-renewing cells capable of differentiating into the wide variety of mature blood lineages (LIN) in our bodies, ranging from red blood cells to the entire spectrum of immune cells^9^ Vertebrate hematopoiesis starts within the yolk sac to succinctly continue over aorta-gonads-mesonephros, placenta and fetal liver (FL), whereby each ontogenic HSC stage is marked by distinct immunophenotypes and transcriptional signatures^10^. The SLAM family members CD48, CD150 and CD244 are differentially expressed among primitive progenitors in adult mouse bone marrow (BM) and currently allow the highest purification grade of LIN^−^/Sca-1^+^/Kit^+^/CD48^−^/CD150^+^/CD41^−^ long-term LT-HSCs^11^. Similarly, highest purity of long-term multilineage reconstituting fetal liver HSCs is obtained from the LIN^−^/Sca-1^+^/Mac-1^+^/CD48^−^/CD150^+^ immunophenotype^12^. The most prevalent HSPC population within FL LIN^−^/Sca-1^+^/Kit^+^ (LSK) during E15.5-E18.5 are Flt3^−^/CD48^+^/CD150^+^ cells comprising megakaryocyte-biased MPP2, a multipotent progenitor (MPP) subset^13^. FL-MPP2 frequencies decline to adult levels around post-natal day 0 (P0), until which these MPP2 show a strong decline in colony formation capacity and in vivo repopulation, a paradigm of a functionally distinct but immunophenotypically identical pre-natal HSPC population as compared to its adult counterpart^14^. FL HSCs have higher regenerative potential than young adult marrow HSCs^15^and predominantly produce myeloid cell outputs in contrast to balanced adult HSCs^16^. However, while the FL HSC phenotype is retained up to the 3^rd^ week of life in mice^17^, neonatal committed myeloid progenitor frequencies do not substantially differ from young adult mice and there is no HSPC immunophenotype specific to newborn mouse BM.

Here we characterized the post-natal HSPC landscape in naked mole-rats, and found a high-frequency myeloid-biased progenitor stage specific to newborn BM and to a lesser extent the spleen. In contrast to murine FL-MPPs the naked mole-rat progenitors concomitantly overexpressed CXCL12 and CXCR7, leading to impaired BM niche homing. Our results provide an unprecedented finding of stage-specific neonatal myeloid progenitors without homing capacity, improving our current understanding of stem cell biology in the naked mole-rat as a model of exceptional longevity.

## Results

### An abundant progenitor population specific to naked mole-rat neonatal hematopoiesis

Previously we functionally characterized purified naked mole-rat HSPC subsets by cross-reactive monoclonal antibodies^6^. Using cross-reactive clones against the human stem cell markers CD34 and THY1 created a complex staining pattern characteristic for adult naked mole-rat bone marrow (BM, Figure 1a). The transcriptional identity of neutrophil granulocytes (GC), erythroid (ERY) and T cells (TC) as well as megakaryocytic erythroid progenitors (MEP) and HSPCs gated from the flow cytometry (FACS) staining was confirmed by CITE-Seq^18^. We then harvested hematopoietic tissues from P2-3 newborns to be profiled by our naked mole-rat HSPC FACS panel. Neonatal BM featured a distinctive Thy1.1/CD34 staining pattern, with less mature GCs than adult BM (Figure 1b). Notably, an abundant neonate Thy1.1^hi^/CD34^hi^ cell population extending from the Thy1.1^int^/CD34^hi^ HSPC fraction was absent in adult animals. A 4.8-fold increase of Thy1.1/CD34 double-positive cells in neonate compared to adult livers indicates fetal hematopoiesis as conserved origin of later developmental steps (Supplementary Figure 1). Reasoning from the Thy1.1^hi^ levels seen exclusively in neutrophils^6^, we referred to Thy1.1^hi^/CD34^hi^ cells as neonate myeloid progenitors (NMP).

**Fig. 1.**
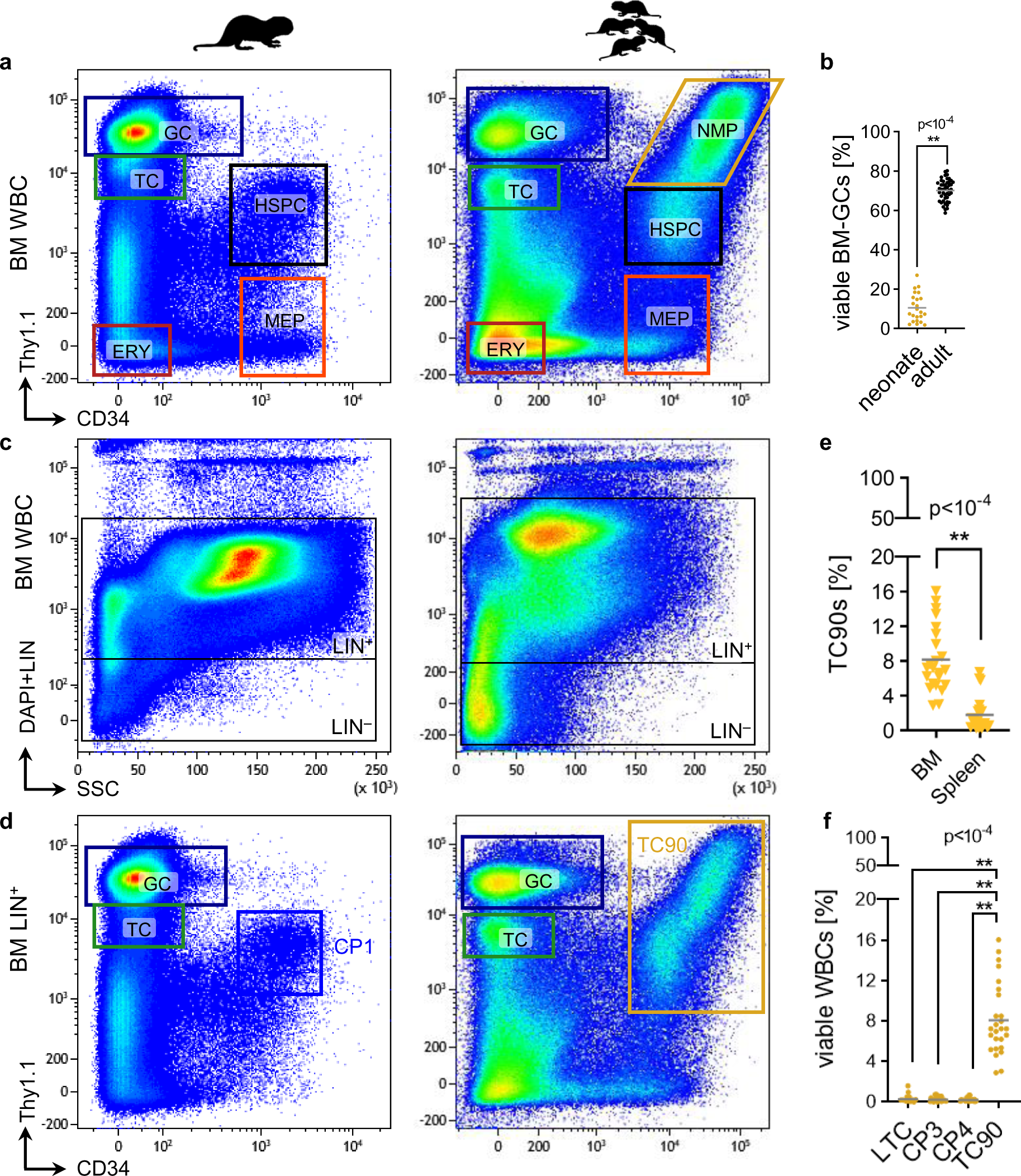
Marrow immunophenotyping reveals a specific neonatal progenitor stage. **a**, Representative FACS gating of viable naked mole-rat adult [left] and neonate [right] whole BM cells. GC, neutrophil granulocytes; TC, T cells; HSPC, hematopoietic stem and progenitor cells; MEP, megakaryocytic erythroid progenitor; ERY, erythroid cells; NMP, neonatal myeloid progenitor. **b**, Frequencies of LIN^+^/Thy1.1^hi^/CD34^−^ GCs in adult (N=46) and neonatal (N=23) BM; p-values were determined by unpaired Student’s t-test. Representative FACS gating of viable naked mole-rat adult [left] and neonate [right] **c**, whole BM or **d**, LIN^+^ BM cells. LIN, lineage cocktail CD11b/CD18/CD90/CD125; TC90, CD11b^−^/CD90^+^/Thy1.1^int/hi^/CD34^hi^. **e**, Frequencies of TC90s in NBM (N=23) and neonatal spleens (N=22); p-values were determined by unpaired Welch’s t-test. **f**, Frequencies of neonatal BM cell fractions. p-value determined by Brown-Forsythe’s One-way ANOVA; N=23.

We previously showed that CD11b and CD18 label naked mole-rat myeloid peripheral blood cells, whereas high CD90-antigen marks neutrophils and dim CD90-antigen T cells^6^ This labelling pattern was confirmed in adult marrow (Supplementary Figure S1). Together with CD125, which labels B cells^7^, we combined a lineage depletion cocktail (LIN) used to further purify the HSPC fraction (Figure 1c). The gating limit for LIN^−^ was positioned to exclude 2 orders of magnitude of positively LIN-stained cells and in parallel to exclude any high complexity side scatter cells. Compared to adult BM neonate marrow featured a higher LIN^−^ fraction, which is expected given the higher amount of erythropoiesis and stem and progenitors required for postnatal development^19^. Specific to neonates however was a strongly expanded LIN^+^/Thy1.1^int/hi^/CD34^hi^ population containing cells in some animals with the highest CD34 and Thy1.1 staining intensities observed in any tissue (Figure 1d). Although this immunophenotype resembled adult LIN^+^/Thy1.1^int^/CD34^hi^ CP1 (<u>c</u>andidate <u>p</u>opulation), neonate LIN^+^/Thy1.1^int/hi^/CD34^hi^ cells expressed CD90-antigen at high levels and did not express CD11b (Supplementary Figure 1). Hence we coined this fraction TC90 (<u>T</u>hy1.1^int/hi^/<u>C</u>D34^hi^/CD<u>90</u>^lo^), which was predominantly found in neonatal marrow at a 4.6-fold higher level compared to spleen (Figure 1e). While there was ∼5-fold increase of CP1 (0.66±0.22%) over LIN^−^/Thy1.1^int^/CD34^hi^ LTCs (0.13±0.07) in adult BM^6^, the NMP population specific to neonate whole BM was exclusively detected in the LIN^+^ TC90 fraction (8.1±3.8%), which comprised 38-fold more cells than neonate LIN^−^ CP2 (0.21±0.21%; Figure 1f).

Next we immunophenotyped murine adult and P3 HSPC fractions, which showed an increased LIN^−^/Sca-1^+^/Kit^+^ (LSK) stem and progenitor compartment in neonates across hematopoietic tissues (Figure 2a). Expansion of LSKs relative to adult tissues was highest in neonatal liver, resembling fetal liver hematopoiesis progressing towards the neonatal-to-adult stage transition (Figure 2b). Cd150^+^ LSK long-term hematopoietic stem cells (LT-HSCs) are slightly elevated in 6 month-old adult marrow, however multipotent progenitors (MPP) are 4.2-fold more abundant in neonatal mice (Supplementary Figure 1). Myeloid progenitor populations within the LIN^−^/Sca-1^−^/Kit^+^ (LK) compartment were unaltered. Naked mole-rat LTCs, which equivalently to murine LSKs have lympho-myelo-erythroid potential and harbor the primitive HSC pool^6^, remain at neonatal levels in adult animals (Figure 2c-d). This is a further neotenic trait, the retention of juvenile features in adult animals, of the naked mole-rat^20^. Consistent with a prevalence of erythropoiesis around birth neonate CP5, comprised of hyperproliferative late erythroblasts^6^, expanded 3.5-fold in BM and 4.7-fold in spleen (Supplementary Figure 2).

**Fig. 2.**
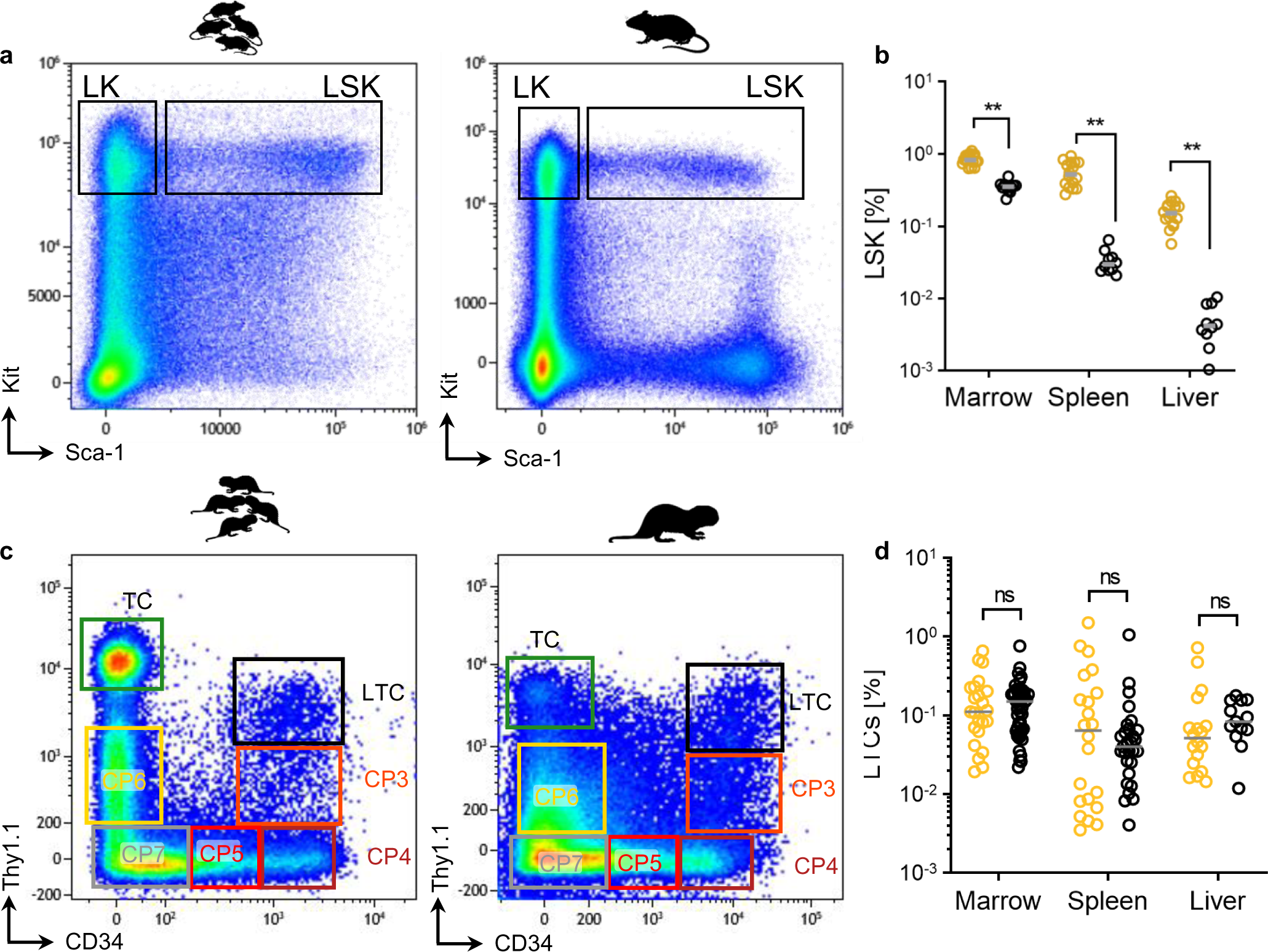
Neonatal HSPC compartment frequencies in mice and naked mole-rats. **a**, Representative FACS gating of viable mouse neonate [left] and adult [right] LIN^−^ BM cells. LK, LIN^−^/Sca-1^−^/Kit^+^; LSK, LIN^−^/Sca-1^+^/Kit^+^. **b**, Frequencies of LSKs across mouse hematopoietic tissues. Adult BM (N=12), neonate BM (N=18), adult spleen (N=10), neonate spleen (N=16), adult liver (N=10), neonate liver (N=16); p-value determined by Sidak’s Two-way ANOVA. **c**, Representative FACS gating of viable naked mole-rat neonate [left] and adult [right] LIN^−^ BM cells. CP3, LIN^−^/Thy1.1^lo^/CD34^hi^; CP4, LIN^−^/Thy1.1^−^/CD34^hi^; CP5, LIN^−^/Thy1.1^−^/CD34^lo^; CP6, LIN^−^/Thy1.1^lo^/CD34^−^; CP7, LIN^−^/Thy1.1^−^/CD34^−^; LTC, LIN^−^/Thy1.1^int^/CD34^hi^. **d**, Frequencies of LTCs across naked mole-rat hematopoietic tissues. Adult BM (N=47), NBM (N=23), adult spleen (N=29), neonate spleen (N=23), adult liver (N=13), neonate liver (N=18); p-value determined by Sidak’s Two-way ANOVA.

Conclusively, we found a large expansion of CD90-antigen positive cells co-expressing naked mole-rat stem cell markers Thy1.1-antigen and CD34 specifically in neonates, whereas primitive stem and progenitor LTCs remained at neonatal levels in adult hematopoietic tissues.

### TC90 cells feature unique transcriptome and splicing signatures

To address molecular changes between neonate and adult HSPCs we integrated RNA-Seq profiles of TC90s in our previously reported dataset from adult CP1/LTC/CP3/CP4^6^. This combined to 9863 expressed orthologues across 24 normalized samples (Supplementary Figure 2). Using t-distributed stochastic neighborhood embedding (t-SNE)^21^ neonate TC90s clearly formed a separate cluster, however adult CP1 and LTCs could not be resolved into their separate immunophenotypes but instead clustered together as CP1/LTC partition, same for CP3/4 (Figure 3a). Differential gene expression analysis found most upregulated genes for TC90s (Figure 3b). Differentially expressed genes (DEGs) were fed into single sample gene set enrichment analysis (ssGSEA) using a curated collection of expression signatures from human, mouse and naked mole-rat HSPCs for cell type annotation^6^ (Supplementary Table 1). The adult partitions CP1/LTC and CP3/4 showed consensus annotation patterns towards their combined FACS population identity: Human HSC, human CD34^+^ and naked mole-rat LTC and lymphomyeloid signatures enriched while naked mole-rat common erythroid signatures depleted for CP1/2; human MEP, naked mole-rat CP4 and common erythroid signatures enriched but lymphoid and myeloid signatures depleted in CP3/4 (Figure 3c). TC90 cells shared a human myeloid gene signature with CP1/2 and were enriched for human and mouse HSC gene expression, resembling a myeloid-biased early progenitor.

**Fig. 3.**
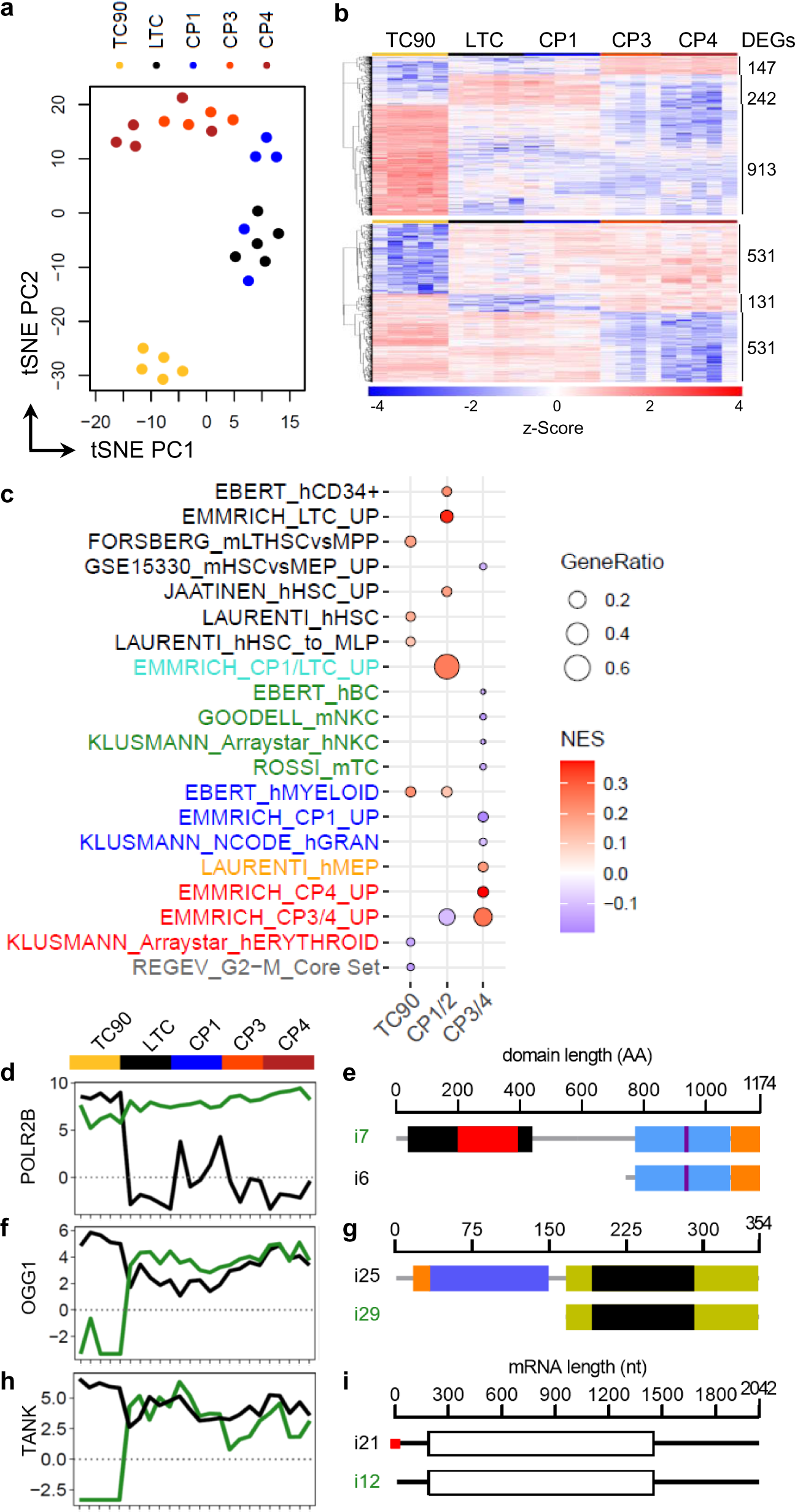
Distinct expression and splicing patterns in neonatal TC90. **a**, Unsupervised t-SNE clustering of neonate TC90 (N=5) and adult LTC (N=5), CP1 (N=5), CP3 (N=4), CP4 (N=5) transcriptomes. LTC/CP1 and CP3/4 form overlapping partitions, TC90s cluster as a distinct group. **b**, Heat- [top panel] and Coldmap [bottom panel] of differentially expressed genes (DEG) for each of the three groupings TC90, CP1/LTC, CP3/4. **c**, GSEA of naked mole-rat neonate and adult BM HSPCs using each DEG signature genes. GeneRatio, (signature ∩ term) / (signature ∩ all terms); NES, normalized enrichment score; displayed the top genesets filtered for GeneRatio > 0.041 and NES > 0.1 | NES < 0.1; each *q* < 10^-4^. Note that the HSC genesets enriched in TC90s are distinct from those enriched in CP1/LTC. **d**, POLR2B; **f**, OGG1; **g**, TANK. Isoform downregulated in TC90, green; Isoform upregulated in TC90, black. **e**, POLR2B domains in amino acids (AA): Rpb2_1 protrusion, black; Rpb2_2 lobe, red; Rpb2_6 hybrid binding & wall, blue; β chain signature, purple; Rbp2_7 anchor & clamp, orange. **g**, OGG1 domains: MTS, orange; OGG_N N-terminal, blue; ENDO3c endonuclease, yellow; HhH-GPD motif, black. **i**, TANK transcripts in nucleotides (nt): TOP motif, red; ORF, white box.

Next we checked whether this profile was also present as differential transcript usage (DTU). Using the same dataset as in Figure 3a we obtained 15038 unique transcripts assigned through the FRAMA BM transcriptome^6, 22^, leading to virtually the same clustering (Supplementary Figure 2). Likewise, TC90 cells had highest DTU, thus we checked for inversely <u>e</u>xpressed transcripts (IETs; see Methods) from the same gene locus in TC90s (Supplementary Figure 2).

Out of 12 total IETs we found an isoform i6 of POLR2B strongly upregulated exclusively in TC90 (1260-fold p<10^-11^; Figure 3d). BLAST alignments of the mRNA coding regions (ORF) revealed an ubiquitously expressed POLR2B isoform i7 downregulated in TC90 (3.6-fold p<10^-6^) giving rise to the full-length RNA-Polymerase-2β subunit protein. The truncated i6 lacks protrusion and lobe domains (Figure 6e), which contribute to maintenance of the transcription bubble but are known as dispensable region in bacterial homologues^23^. Another IET pair was found at the OGG1 locus, a DNA repair enzyme excising 8-oxoguanine which can be formed upon reactive oxygen species (ROS) exposure^24^. TC90s overexpressed the i25 full-length isoform (5.5-fold p<10^-7^), while expression of a truncated i29 lacking both the mitochondrial targeting signal and an N-terminal domain contributing to substrate recognition was completely abolished (Figure 3f-g). The IET isoform with the strongest depletion for TC90s was i12 of the TANK gene (890-fold, p<10^-4^; Figure 3h), an adaptor binding to TRAF1-3 and thereby modulating TNFR:NF-κB signaling^25^. Interestingly, the two isoforms solely differed in their 5’UTR, with only the i21 isoform harboring a 9bp TOP motif associated with mTORC1-mediated translational induction^26^ (Figure 3i).

Overall neonate TC90s differ from adult naked mole-rat LTCs and progenitors by very high abundances in spleen and BM, high CD90-antigen levels and specific alternative splicing patterns.

### The SDF-1α/CXCR4-axis is altered in TC90 cells

The ontogeny-associated differences between murine fetal and adult hematopoietic tissues are conserved in humans^27^. To segregate the inherent properties of TC90s as an earlier progenitor pool in naked mole-rats, we compared them to LTCs along with human cord blood (CB) and marrow HSCs (Figure 4a, Supplementary Figure 3). Using the MSigDB hallmark geneset collection^28^we classified the stem cell types according to their signatures, which yielded Epithelial-Mesenchymal-Transition (EMT) as top ssGSEA category for TC90 cells (Figure 4b). Importantly, the top leading edge gene for EMT in TC90s was CXCL12 (Figure 4c). To our surprise we found the cognate receptor CXCR4 as one of the top downregulated genes in TC90s. Apart from its primary target CXCL12 can also bind to the alternate chemokine receptor homolog ACKR3^29^, leading to ligand internalization and β-arrestin recruitment. Strikingly, TC90 cells specifically overexpressed all three ACKR interceptors (Figure 4c). The arrestin family of adaptor proteins can induce MAPKs, AKT but also NF-κB signaling^30^, the latter could be further modulated by i21 TANK activation (Figure 3h-i). Consistent with a fetal myeloid predominance TC90s overexpressed CD14 and CD38, along with the stem cell master transcription factor EBF1 (Figure 4c). On the contrary there are key marker genes of the naked mole-rat stem cell state emerging from population and single cell RNA-Seq such as ICAM2, HOXA9, GATA2, and HHEX, MEP markers MDK, NFE2 and conserved HSC regulators TAL1, BMI1, MEIS1 and HMGA2, appear strongly decreased in TC90s (Figure 4c, Supplementary Table 2). An intriguing finding are the coordinated upregulation of antiproliferative p21, BTG1 and 4 IGF-binding proteins together with corresponding reduction of 4 centromer proteins, CEP85, POLR1A and MYB. Despite elevated frequencies of TC90s in neonatal BM, spleen and liver but in line with a depleted G2M-Core Set cell cycle signature (Figure 3c), this suggests the neonate-specific LIN^+^/Thy1.1^hi^/CD34^hi^ extension of the CP1-like LIN^+^/Thy1.1^int^/CD34^hi^ subset within TC90 cells as a granulocytic precursor with little self-renewal and proliferative activity.

**Fig. 4.**
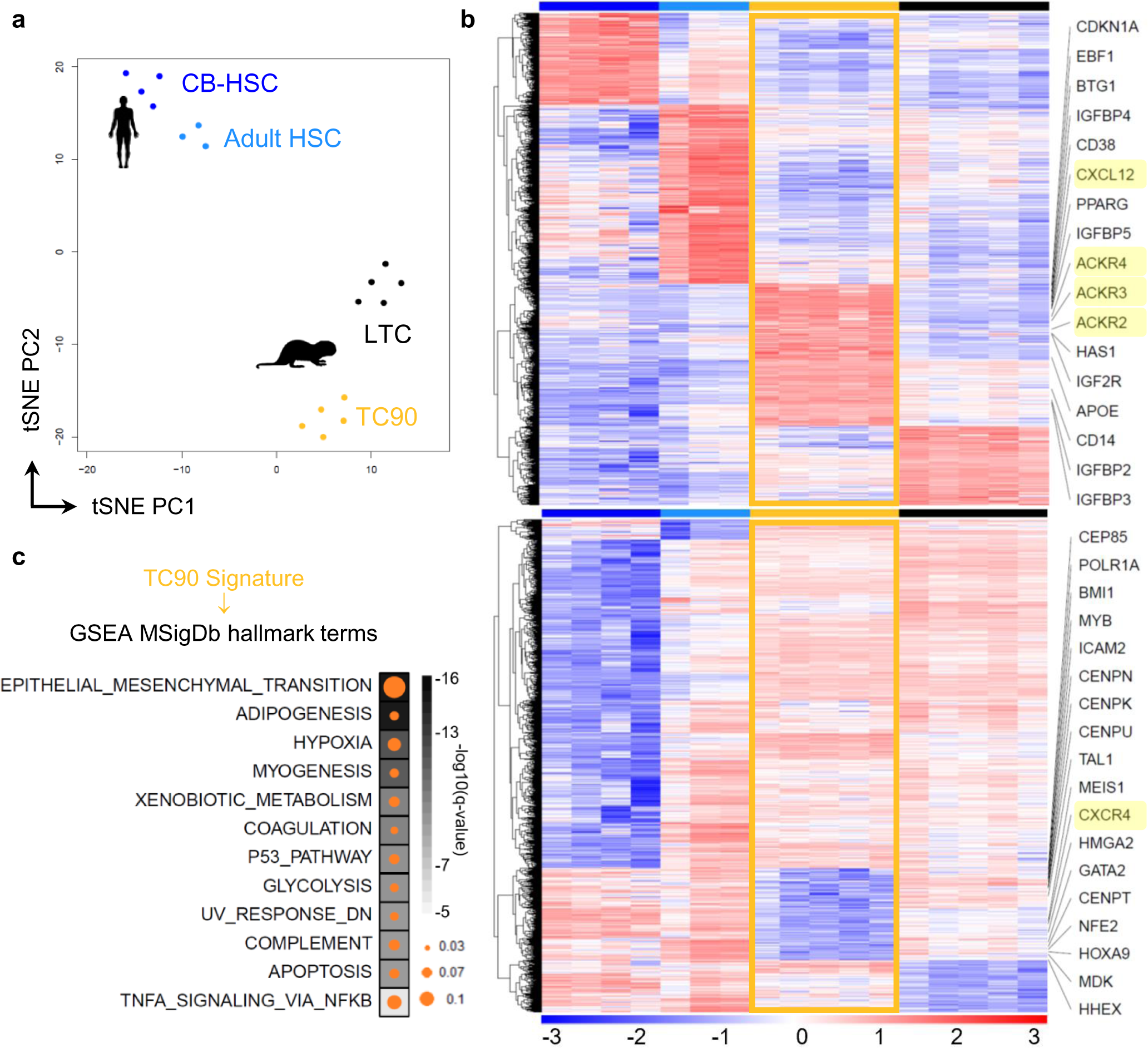
TC90 cells inversely expressed the CXCR4/CXCL12 ratio. **a**, Unsupervised t-SNE clustering, color code used on heatmaps throughout this Figure. CB-HSC, human neonatal cord-blood HSCs; Adult HSC, human adult HSC; LTC, adult naked mole-rat HSPCs. **b**, Heatmap showing upregulated [top] and downregulated [bottom] orthologs with framed TC90 DEG signature; z-score derived from voom-transformed transcripts per million (TPMs). **c**, GSEA of TC90 DEG signature displaying the top 10 q-value terms filtered for NES > 0.1 and GeneRatio > 0.02; each q < 10^-6^. Circle size, GeneRatio; Square color, -log10(*q*-value).

Whereas an absence of HSC maintenance genes would be expected in a committed progenitor such as TC90, we aimed to confirm the inversed CXCR4/ACKR expression pattern by extending the comparison to human HSCs and committed lymphomyeloid progenitors. Thus we integrated expression profiles of common lymphoid progenitors (CLP), multipotent lymphoid progenitors (MLP), granulocytic monocytic progenitors (GMP) and HSCs from 6 different RNA-Seq datasets^31–35^ across cord blood and adult BM stages with LTC, CP1 and TC90 transcriptomes (Figure 5a). This compound dataset comprised 51 samples and 10291 common orthologs (Supplementary Figure 3). Unsupervised hierarchical clustering faithfully recapitulated each of the two ontogenic stages and species groupings (Figure 5b). Remarkably, CXCL12 and ACKR3, also known as CXCR7, were specifically overexpressed in TC90s as compared to all other cell types (Figure 3c). Concordantly, CXCR4 levels were lowest in naked mole-rat TC90 cells. While ACKR2 was also overexpressed in TC90, the alternate CXCR7 ligand CXCL11 was absent in naked mole-rat HSPCs (Supplementary Figure 3). Interestingly, HMGA2 which maintains the fetal stem cell status in concert with LIN28B^36^, was higher expressed throughout human neonatal stem and progenitor populations, whereas naked mole-rat TC90 showed less HMGA2 transcription than adult LTCs and CP1 (Supplementary Figure 3). The same pattern was seen for MLLT3, the fusion partner AF9 of t(9;11) MLL-rearranged translocations, which has a role in early mouse hematopoiesis^10, 37^, whereas KMT2A was generally lower expressed in naked mole-rats and MLLT1 showed no significant changes across species, cell types or ontogenies (Supplementary Figure 3). The ssGSEA profile of TC90 cells compared to neonate and adult HSPCs reproduced the EMT signature as top MSigDb hallmark term (*q*<10^-29^; Supplementary Table 3). Using the fGSEA tool to retrieve the leading edge genes also scored EMT as top term (*q*=0.00055; Supplementary Table 3), which were enriched with multiple secreted proteins responsible for organization of or interacting with the extracellular matrix (ECM), such as Collagen Type I Alpha 2 Chain, PCOLCE, Fibronectin 1, Fibulin 1/2/5, Elastin, Transgelin, Thrombospondin 1, Syndecan 1/4, Periostin, Nidogen 2, Versican, Decorin, ECM1 and MGP (Figure 5d). In conjunction with elevated skeletal muscle-related TPM1, GEM, SGCB transcripts TC90s overexpressed several factors implicated in cell migration and motility. Strikingly, THY1 mRNA expression across species and ontogenic stage correlated with the TC90 Thy1.1^hi^ subset immunophenotype (Figure 1d, 5d).

**Fig. 5.**
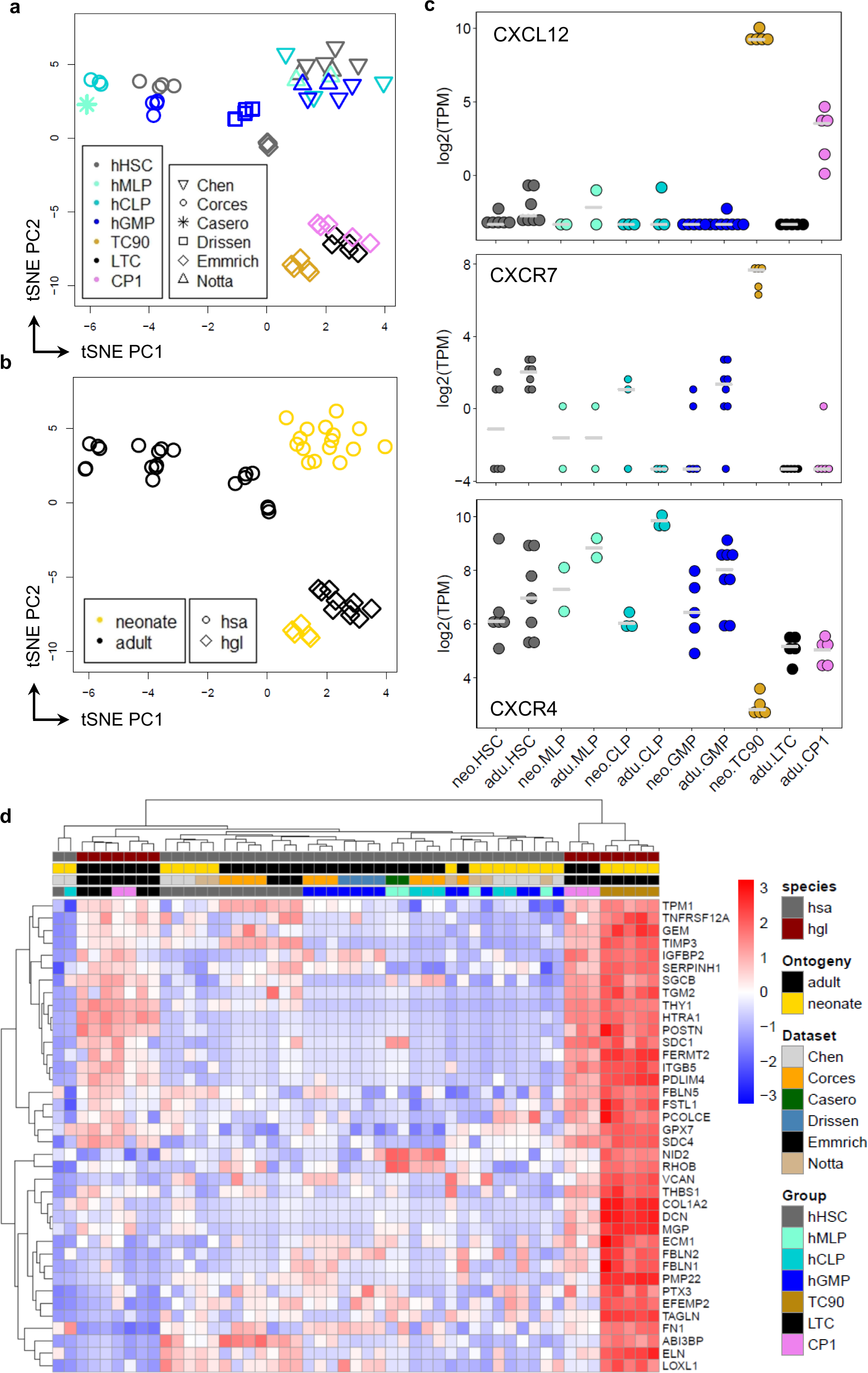
Extended cross-species comparison confirms CXCR7/CXCL12 overexpression. Unsupervised t-SNE clustering with **a**, Colors for cell types, symbols for datasets; **b**, Colors for ontogeny, symbols for species. hHSC, human HSCs; hMLP, human multipotent lymphoid progenitor; hCLP, human common lymphoid progenitor; hGMP, human granulocytic monocytic progenitor; hsa, *homo sapiens*; hgl, *heterocephalus glaber*. **c**, Log-transformed TPM expression across cell types, species and ontogenies for orthologs CXCL12 [top], CXCR7 [middle] and CXCR4 [bottom]. **d**, Heatmap the leading edge genes of the top GSEA term EPITHELIAL_MESENCHYMAL_TRANSITION (*q* < 5.5×10^-4^; NES = 2.54); z-score derived from voom-transformed TPMs.

**Fig. 6.**
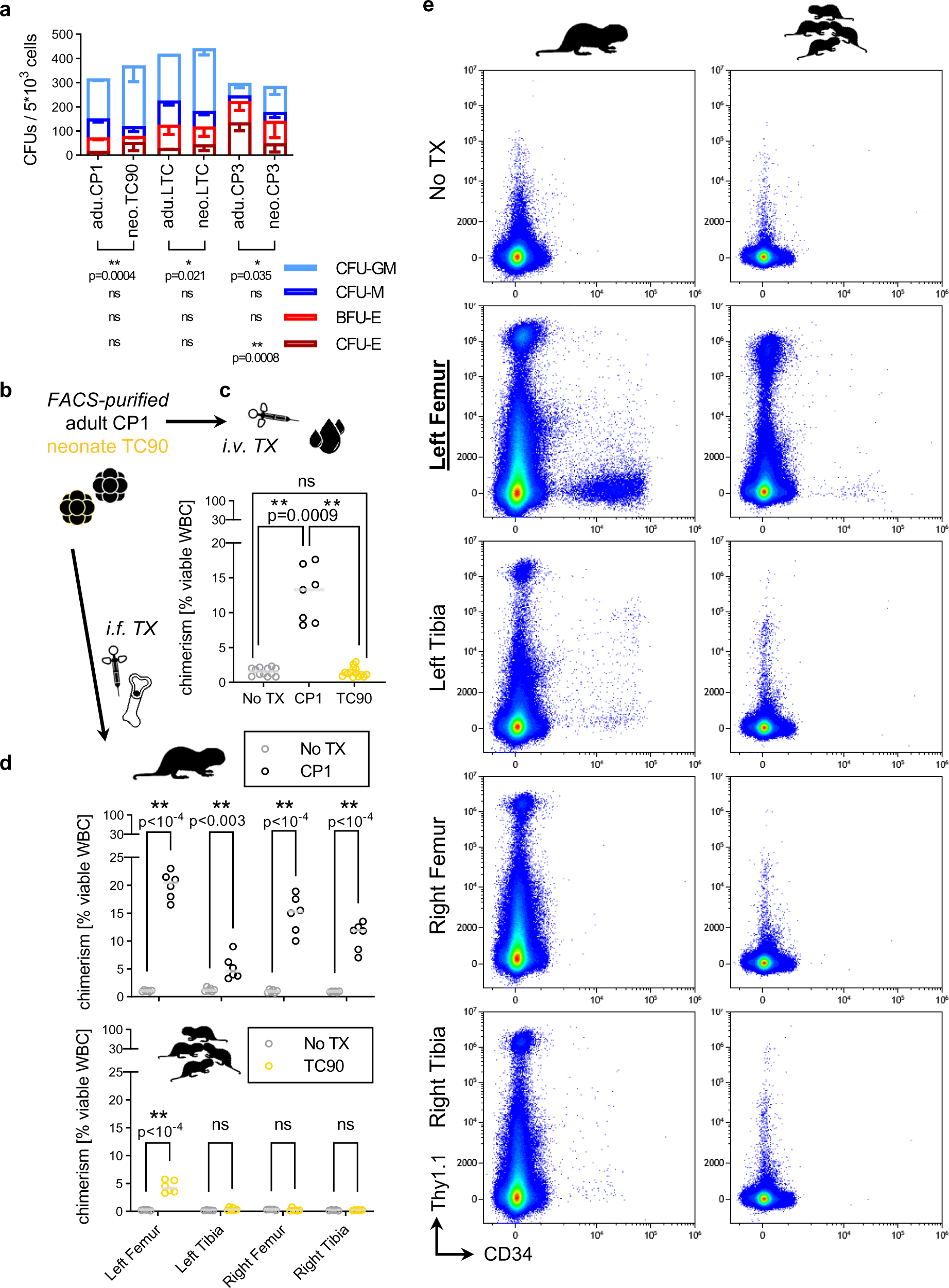
TC90s are myeloid-biased progenitors without homing capacity *in vivo*. **a**, Colony assay of sorted naked mole-rat adult (N=4) and neonate (N=3) BM HSPCs, grown at 32°C for 3 weeks. Error bars denote s.d., p-value determined by Two-way ANOVA, corrected for multiple testing with Two-stage linear step-up procedure of Benjamini, Krieger and Yekutieli. **b**, Naked mole-rat adult CP1 or neonate TC90 cells were FACS-purified and xenotransplanted (TX) into NSGS hosts via **c**, Intravenous (*i.v.*) injection, host whole BM from all combined long bones at 4 weeks post TX. Chimerism gated as viable cells positive for either or both naked mole-rat cross-reactive HSPC markers CD34 and Thy1.1-antigen. **d**, Intrafemoral (*i.f.*) injection, host whole BM from each single long bone at 4 weeks post TX. e, Representative FACS gating for chimerism on *i.f.* injected hosts with adult CP1 [left panel] or neonate TC90 [right panel].

Here we report the first developmental stage-specific gene expression pattern for naked mole-rat neonatal committed progenitor TC90 cells, marked by a conserved EMT signature, CXCL12 signaling through ACKRs and absence of multiple HSC maintenance genes. Moreover, high expression of a broad ECM component spectrum suggests a role in remodeling of the bone marrow microenvironment.

### Neonate TC90 are myeloid-biased CMP-like progenitors without homing capacity

The unique transcriptome signature of neonatal TC90 was further interrogated using colony formation and xenotransplantations (TX) of sorted naked mole-rat HSPCs. Multipotency as the capacity to form multiple distinct lineages via differentiation can be assayed through quantitation of progenitor frequencies from colony assays in methylcellulose^38^. We previously validated the scoring of colony-forming unit (CFU) types by cytochemistry of single colonies^6^. Here we show representative morphologies for each colony type derived from naked mole-rat adult marrow LTCs (Supplementary Figure 2). CFU-M (<u>M</u>onocytic/Macrophage) colonies displayed colorless, large cells as scattered aggregations, CFU-GMs (<u>G</u>ranulocytic-<u>M</u>onocytic) had high amounts of scattered single neutrophils, whereas BFU-Es (<u>B</u>urst-forming <u>u</u>nit <u>E</u>rythroid) were larger clusters of condensed small cells with brownish coloration indicating heme production, and CFU-Es (<u>Colony</u>-forming <u>u</u>nit <u>E</u>rythroid) had smaller clusters of condensed very small cells with brown-to-red coloration. Upon comparison between neonate and adult sorted HSPC fractions we found that all neonate populations produced significantly more CFU-GM (<u>G</u>ranulocytic-<u>M</u>onocytic) colonies (Figure 6a). Neonatal CFU-GM colonies were generally of larger diameters than any other colony type (Supplementary Figure 4). Given the myeloid bias of ontogenetically younger fetal liver cells in mice this was expected^16^. Moreover, we saw neonate CP3 MEPs yielding less CFU-E colonies (2.8-fold, p=0.0017), which derive from more differentiated erythroid precursors than BFU-E clones, indicating a higher frequency of early progenitors within the erythroid lineage pool of neonate MEPs.

When adult LTCs or CP1 were transplanted via intravenous injection (i.v.) into immunodeficient mice (NSGS), both produced similar peak engraftment rates at 12-16% host BM chimerism between 2 and 4 weeks post TX^6^. Thus we compared CP1 with neonate TC90 xenografts administered either i.v or through intrafemoral injection (i.f.) to account for potential engraftment failures due to impaired BM homing (Figure 6b). At 4 weeks post TX through i.v. there was clearly no engraftment of TC90s, whereas adult CP1 reached 12.6±4% BM chimerism (Figure 6c). Next we quantified i.f. xenografts by separating each femur and tibia from recipients to distinguish the direct marrow injection site of the left femur from the left tibia and contralateral bones (Figure 6d-e). Adult CP1 produced robust multilineage engraftment of 20±2.4% at the injected femur. Reconstitution of non-injected bones requires HSPC homing through the circulation, resulting in diminished engraftment distal the i.f. injection site^39^. Conversely, CP1 xenografts displayed decreasing chimerism in contralateral right femori with 14.8±3.3%, right tibiae with 10.9±2.5% and left tibiae with 5.3±2.1%. Strikingly however, TC90 chimerism yielded 4.4±1.1% at the i.f. site, whereas non-injected bones had no engraftment (Figure 6d). Conclusively, HSPC homing in neonate TC90 is abrogated through downregulated CXCR4 expression accompanied by high ACKR levels acting as decoys for SDF-1α signaling. Previously we applied the Thy1.1/CD34 staining pattern to differentiate between the major xenograft lineages^6^. The TC90 grafts of left femori have very little CD34^+^ engraftment compared to CP1 (Figure 6e). Instead Thy1.1^hi^/CD34^−^ GCs form the bulk of TC90 progeny, which is in line with their high frequency of CFU-GM colony formation and naked mole-rat neutrophilic Thy1.1-antigen and CD90-antigen levels at the cell surface. Thus we showed that TC90 cells have common myeloid progenitor (CMP) potential with a strong bias towards neutrophil cell fate and no capacity of HSPC homing.

## Discussion

Naked mole-rats are the longest-lived rodents that remain healthy until the end of their lives and are resistant to age-related diseases including cancer. Fetal and neonatal stem cells are programmed in a developmental stage-specific manner to establish hematopoiesis and become the progeny of the adult HSC pool, which is crucial for lifelong blood and immune cell production. Thus characterization of neonatal HSPCs will improve understanding of the biology that enables the exceptional lifespan of naked mole-rats.

Here we show that naked mole-rat neonatal BM is enriched with a specific progenitor stage we termed TC90. Mice showed a substantial expansion of neonatal MPPs responsible for higher LSK HSPC frequencies throughout hematopoietic tissues, while their more committed progenitors remained unchanged towards the adult stage. By contrast, the primitive HSPC compartment marked by LTCs in naked mole-rats remained unchanged, while TC90s are expanded over their adult counterpart CP1, which is a myeloid-biased CMP-like progenitor with durable self-renewal as shown by CITE-Seq and FACS^6^. Moreover, the majority of TC90 cells are CD11b^−^/CD90^hi^/Thy1.1^hi^/CD34^hi^, according to Thy1.1-antigen levels they extent from the CP1 LIN^+^/Thy1.1^int^/CD34^hi^ immunophenotype, thereby forming an intermediate differentiation stage likely linked to CD11b^+^/CD90^hi^/Thy1.1^hi^/CD34^−^ mature GCs. Whole BM of naked mole-rats comprises ∼74% neutrophil GCs and their precursors, whereas mice have ∼52% and humans ∼61% GCs^6, 40^. Hence the accumulation of a neonatal myeloid progenitor (Figure 1a), dedicated to increase innate immune effector GCs, appears to be an adaptation to protect the newborn and sustain ample neutrophil frequencies in adult marrow and blood. It is important to note that the TC90 sorting and quantitation gate subsumes a heterogenous population of neonatal CP1 and committed granulocytic precursors, based on the CD11b^−^/Thy1.1^hi^/CD90^hi^ NMP immunophenotype (Figure 1a). This explains the shift towards granulocytic differentiation and accounts for the decreased erythroid commitment seen for TC90 in colony assays and i.f. xenografts.

TC90 RNA-Seq profiles revealed downregulation of CXCR4 and concurrent overexpression of several ACKR orthologs and CXCL12, suggesting alternate SDF-1α/CXCR4-axis signaling. Indeed TC90s produced xenografts upon direct marrow transfer at the injection site, yet host marrow distal from the i.f. site was not repopulated, and i.v. delivered transplants failed to engraft. This indicates a BM homing deficiency in TC90 due to competitive inhibition of CXCR4 by CXCR7^41^. CXCR7 can heterodimerize with CXCR4 and interfere with CXCR4-mediated integrin activation but not chemotaxis^42, 43^. CXCL11 triggers CXCR7-induced cell adhesion in myeloid leukemia cells^44^. In T cells CXCR7/CXCL12 interaction promotes survival and chemotaxis in a G-protein independent manner^45^, instead signaling occurs through β-arrestins^46^. Conceivably, their exceptionally high transcript levels render TC90 cells as a potential key source of CXCL12 in naked mole-rat neonatal BM. Furthermore, TC90 overexpress multiple secreted factors for ECM organization, pointing towards a role in modeling of the bone marrow niche. If responsible for generating the chemotactic SDF-1α gradient, TC90 cells employing CXCR7 to counteract CXCR4-mediated migratory signaling could remain in and contribute to the developing niche. This would explain both the TC90 insensitivity to homing signals in a transplantation setting, as well as TC90 frequencies of 8.1±3.8% in neonatal whole BM (Figure 1f), enabling sustained production of ECM components. An hematopoietic progenitor stage with native impaired BM homing acting as source of multiple secreted niche factors has not been described in mouse or humans, and might reflect a combination of habitual adaptations to subterranean challenges of naked mole-rat immune systems, slower rate of postnatal maturation^47^, and an altered ECM enriched with high molecular weight hyaluronic acid^48^.

In conclusion, we found an abundant neonatal myeloid progenitor stage that supplies the newborn with an essential reservoir of neutrophils to support a myeloid-based immune system and potentially acts as source of the SDF-1α homing chemokine and multiple ECM components.

This reveals novel aspects of myelopoiesis in the developing blood system of naked mole-rats and outlines transient developmental stages promoting neutrophil accumulation and producing CXCL12 and CXCR7 as unique traits of these long-lived rodents.

## Materials and Methods

### Animals

All animal experiments were approved and performed in accordance with guidelines instructed by the University of Rochester Committee on Animal Resources with protocol numbers 2009-054 (naked mole rat) and 2017-033 (mouse). Naked mole rats were from the University of Rochester colonies, housing conditions as described^1^. C57BL/6 mice were obtained from NIA, immunodeficient NSGS [NOD.Cg-*Prkdc^scid^ Il2rg^tm1Wjl^* Tg(CMV-IL3,CSF2,KITLG) 1Eav/MloySzJ] were purchased from JAX.

### Primary cell isolation

Marrow from neonatal and adult mice and naked mole-rats was extracted from femora, tibiae, humeri, iliaci and vertebrae by crushing. Spleen, liver, thymus and lymph nodes were minced over a 70µm strainer and resuspended in FACS buffer.

### Methylcellulose colony assays

Fresh sorted or whole BM naked mole-rat cells were tested to grow in mouse (M3434, SCT), rat (R3774, SCT) or human (H4435, SCT or HSC005, RnD Systems) methylcellulose formulations to show the highest colony numbers, colony sizes and cell viability with human cytokine cocktails. Unless otherwise stated, 2×10^3^ sorted naked mole-rat cells were added to 3ml of HSC005 supplemented with 1% Penicillin/Streptomycin and 1X GlutaMAX (both Thermo Fisher), grown for 14d at 32°C, 5% CO_2_ and 3% O_2_, and scored.

### Flow Cytometry

<u>Flow cytometry analysis</u> was performed at the URMC Flow Core on a LSR II or LSRFortessa (both BD), or a CytoFlex S (Beckman Coulter). Kaluza 2.1 (Beckman Coulter) was used for data analysis. Staining and measurement were done using standard protocols. Red blood cell lysis was done by resuspending marrow pellets in 4ml, spleen pellets in 1ml and up to 500µl blood in 20ml of RBC lysis buffer, prepared by dissolving 4.1g NH_4_Cl and 0.5g KHCO_3_^−^ into 500ml double-distilled H_2_O and adding 200µl 0.5M EDTA. Marrow and spleen were incubated for 2min on ice, blood was lysed for 30min at room temperature. Cells were resuspended in FACS buffer (DPBS, 2mM EDTA, 2% FBS [Gibco]) at 1×10^7^ cells/ml, antibodies were added at 1µl/10^7^ cells, vortex-mixed and incubated for 30min at 4°C in the dark. DAPI (Thermo Fisher) @ 1µg/ml was used as viability stain. The primary gating path for all unfixed samples was: scatter-gated WBC (FSC-A vs SSC-A) => singlets1 (SSC-W vs SSC-H) => singlets2 (FSC-W vs FSC-H) => viable cells (SSC vs DAPI) == proceed with specific markers/probes. Compensation was performed using fluorescence minus one (FMO) controls for each described panel.

<u>Immunophenotyping</u> of naked mole-rat BM, spleen, liver: CD90 FITC; CD125 PE; Thy1.1 PE-Cy7; CD34 APC, CD11b APC-Cy7. Quantification of murine BM SLAM HSCs was performed using mouse LIN Pacific Blue; Sca-1 BUV395; CD150 PE; Kit PE-Cy7; CD48 APC-Cy7. Quantification of human BM LT-HSCs was performed using human LIN Pacific Blue; CD34 APC; CD38 APC-Cy7; CD45RA FITC; CD90 PE-Cy7. Fluorescence minus one (FMO) controls were applied for fluorescent spillover compensations for each species and tissue used.

<u>Sorting</u> was performed at the URMC Flow Core on a FACSAria (BD) using a 85μm nozzle, staining was done as described. Naked mole-rat HSPC populations were sorted as described with a lineage cocktail comprised of CD11b, CD18, CD90 and CD125 (NMR LIN). Naked mole-rat marrow sorting panel was: NMR LIN Pacific Blue; Thy1.1 PE-Cy7; CD34 APC.

### Xenotransplantations

Naked mole-rat BM and/or spleen cells were extracted, sorted and directly transplanted into 2.5Gy-irradiated (24h pre Tx) NSGS recipients between 5-9 weeks of age at cell doses between 50-200k cells. Injections were done via the retroorbital sinus (i.v.) or through the left femur (i.f.). Hosts were culled at 4 weeks and engraftment frequencies were estimated by flow cytometry using only naked mole-rat markers not cross-reactive with mouse cells and CD45.1 (A20, BioLegend). Engraftment rates were adjusted for input cell dose to 100k/Tx. Intrafemural injections were analyzed by marrow extraction from each hind limb long bone separately.

### Population RNA-Seq

RNA from sorted human and naked mole-rat populations was sequenced at ∼100 million reads on a HiSeq2500v4 (Illumina), the SMARTer® Ultra® Low RNA Kit (Takara) was used for library preparation. All GEO datasets for human and mouse HSPC populations were acquired with SRA toolkit and processed from raw fastq files. Raw Illumina paired-end sequencing reads where subjected to base quality trimming using Trimmomatic^49^ and were assessed with FastQC. *RSEM v1.3.0* with *STAR* aligner option was used to calculate expected counts and TPMs^50^. We used a customized perl script to run RSEM with the FRAMA transcriptome as reference using *bowtie2* aligner option. We also run RSEM with the ENSEMBL94 hetgla_female_1.0 annotation using *STAR* aligner option to confirm all clusterings and differential gene expression signatures for all naked mole-rat samples, results were almost identical to those obtained with FRAMA (data not shown).

Subsequent analysis was done with *R 4.0.2* and *Bioconductor*^51^. Expected counts from different transcript isoforms of the same gene were added up to one unique identifier (uniquefy) using `ddply` and `numcolwise` functions of the *plyr* package, *edgeR* was used to calculate size factors with `method=”RLE”` and computing CPMs. We applied *genefilter* to calculate the interquartile range (IQR) of CPMs with `IQR(x) > 1` to filter unexpressed and outlier genes; library-size normalized, IQR-filtered log2-transformed CPMs were vst-transformed by *DESeq2*, then a PCA from stats package was used as input for *Rtsne*. We applied *limma* to perform voom-transformation and select for differentially expressed genes (DEGs) with p < 0.05 and log-fold-change 1.

GSEA was performed using the *gsva* package with `method=”ssGSEA”` using either the hematopoietic stem and progenitor geneset collection modified from Schwarzer & Emmrich et al^52^ used previously or the MSigDB v6.0 hallmark genesets with a p-value threshold of 0.05^53–55^. All GSEA calculations were performed on the combined up- and downregulated DEG signature for each group, see Supplementary Table 1. The *fGSEA* package was used to retrieve leading edge genes after reperforming GSEA under default conditions^56^, the required gene rank metric was generated after Plaisier et al^57^. All naked mole-rat population RNA-Seq DEG signatures were used to create genesets and were added to Table S1.

The IET (Inversely Expressed Transcript) pairs shown in Figure 3 have been determined by intersecting gene loci of 1197 up- with loci from 846 downregulated isoforms. Pfam and Expasy ID links for domains retrieved by using NCBI Search for Conserved Domains^58^ on the respective FRAMA transcripts were <u>pfam04563</u>: RNA_pol_Rpb2_1; <u>pfam04561</u>: RNA_pol_Rpb2_2; <u>pfam00562</u>: RNA_pol_Rpb2_6; RNA_POL_BETA <u>PS01166</u> expasy signature; <u>pfam04560</u>: RNA_pol_Rpb2_7; <u>cd00056</u>: ENDO3c; <u>pfam00730</u>: HhH-GPD; <u>pfam07934</u>: OGG_N; putative MTS taken from^24^; 5’TOP motif detected by RegRNA 2.0^59^.

For the 2-species comparison in Figure 4 CB-HSC samples were taken from the BLUEPRINT dataset,^32^ adult BM-HSCs and naked mole-rat LTC taken from our hematlas^6^, and TC90 were from this study; For the 2-species comparison uniquefied human and naked mole-rat TPM datasets (Figure 4a, 5a-b) were merged based on HGNC symbols, then *genefilter* was used to calculate IQR of TPMs with `IQR(x) > 1` to filter unexpressed and outlier genes; The *TCC* package was used to calculate TMM-based size-factors. The function `betweenLaneNormalization` with median scaling from the *EDASeq* package was used to normalize for sequencing batch effects. TPMs were vst-transformed by *DESeq2*, then a PCA from the *stats* package was used as input for *Rtsne*; DGE and GSEA were performed as above using the population species as contrast and the MSigDB v7.0 hallmark genesets.

### Quantification and Statistical Analysis

Data are presented as the mean ± SD. Statistical tests performed can be found in the figure legends. P values of less than 0.05 were considered statistically significant. Statistical analyses were carried out using Prism 9 software (GraphPad) unless otherwise stated.

## Supplementary Material

Supplementary Tables can be found in the online version of this manuscript.

## Data availability statement

The RNA-Sequencing data that support the findings of this study are available in figshare.com with the identifiers <u>10.6084/m9.figshare.c.5472735, 10.6084/m9.figshare.c.5472684, 10.6084/m9.figshare.16734475.v1</u>.

## Acknowledgements

The authors thank the Center for Integrated Research Computing (CIRC) at the University of Rochester for providing computational resources and technical support and the URMC Flow Core for assistance with sorting. This work was supported by the US National Institutes of Health grants to V.G. and A.S. S.E. is a fellow of HFSP.

## Author Contributions

S.E. designed and supervised research, performed experiments and analyzed data; S.E., A.S. and V.G. wrote the manuscript.

## Supplementary Figures and Legends

**Supplementary Fig. 1.**
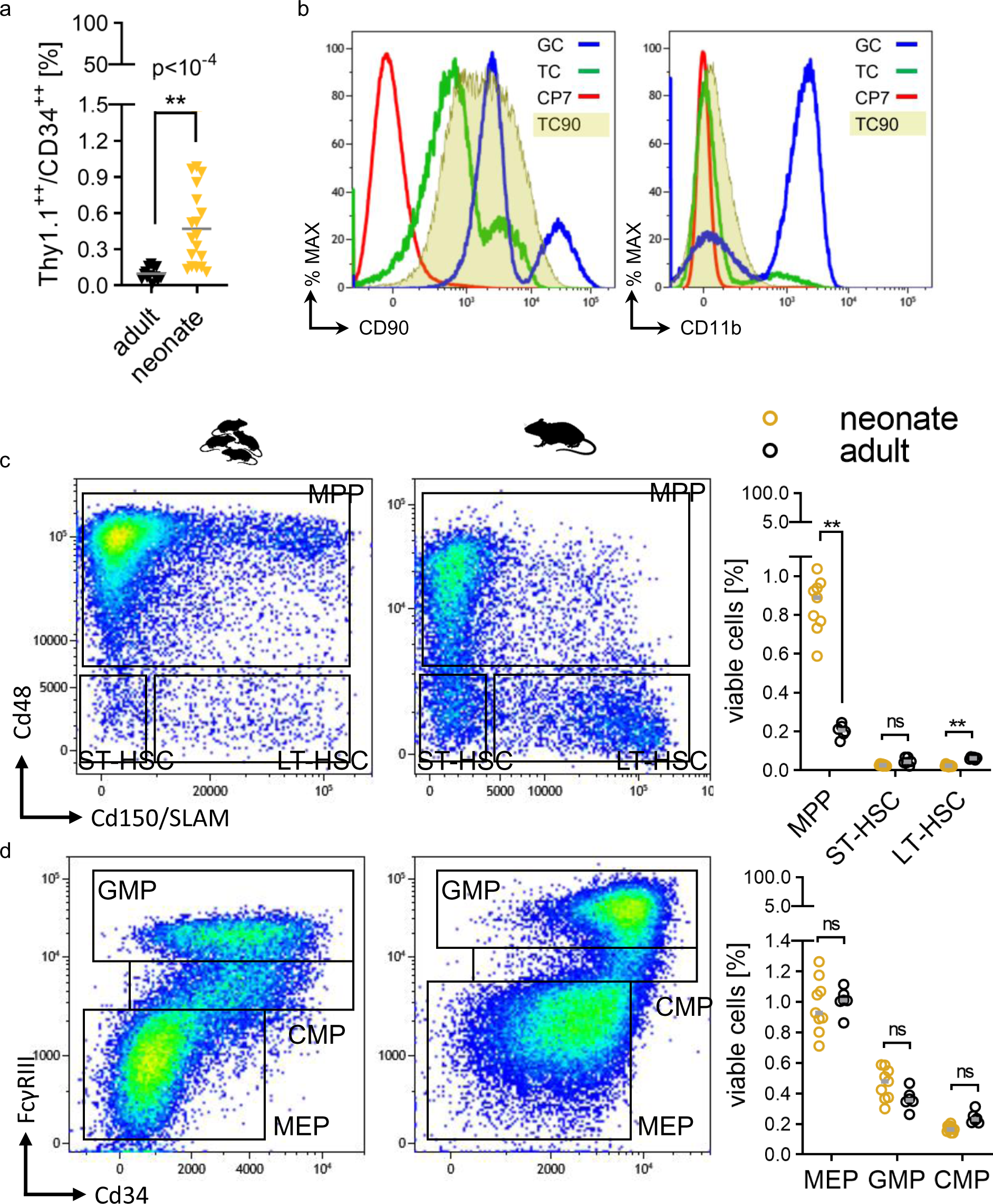
TC90 immunophenotyping, Mouse stem and progenitor FACS. **a**, Frequencies of Thy1.1^+^/CD34^+^ cells in adult (N=13) and neonatal (N=18) livers; p-values were determined by unpaired Welch’s t-test. **b**, Single color flow cytometry histograms of neonatal BM co-stained for CD90-antigen [left panel] and CD11b [right panel] and the LIN/Thy1.1/CD34 labelling; blue line, LIN^+^/Thy1.1^hi^/CD34^−^ GCs; green line, Thy1.1^int^/CD34^−^ TCs; red line, LIN^−^/Thy1.1^−^/CD34^−^ CP7 as negative controls; shaded AUC, LIN^+^/Thy1.1^int/hi^/CD34^hi^ TC90s. Representative FACS gating of neonate [left panel] or adult [right panel] BM **c**, LSKs or **d**, LKs; Frequencies of adult (N=6) and neonatal (N=9) populations see scatter respective plots, p-values were determined by Sidak’s Two-way ANOVA.

**Supplementary Fig. 2.**
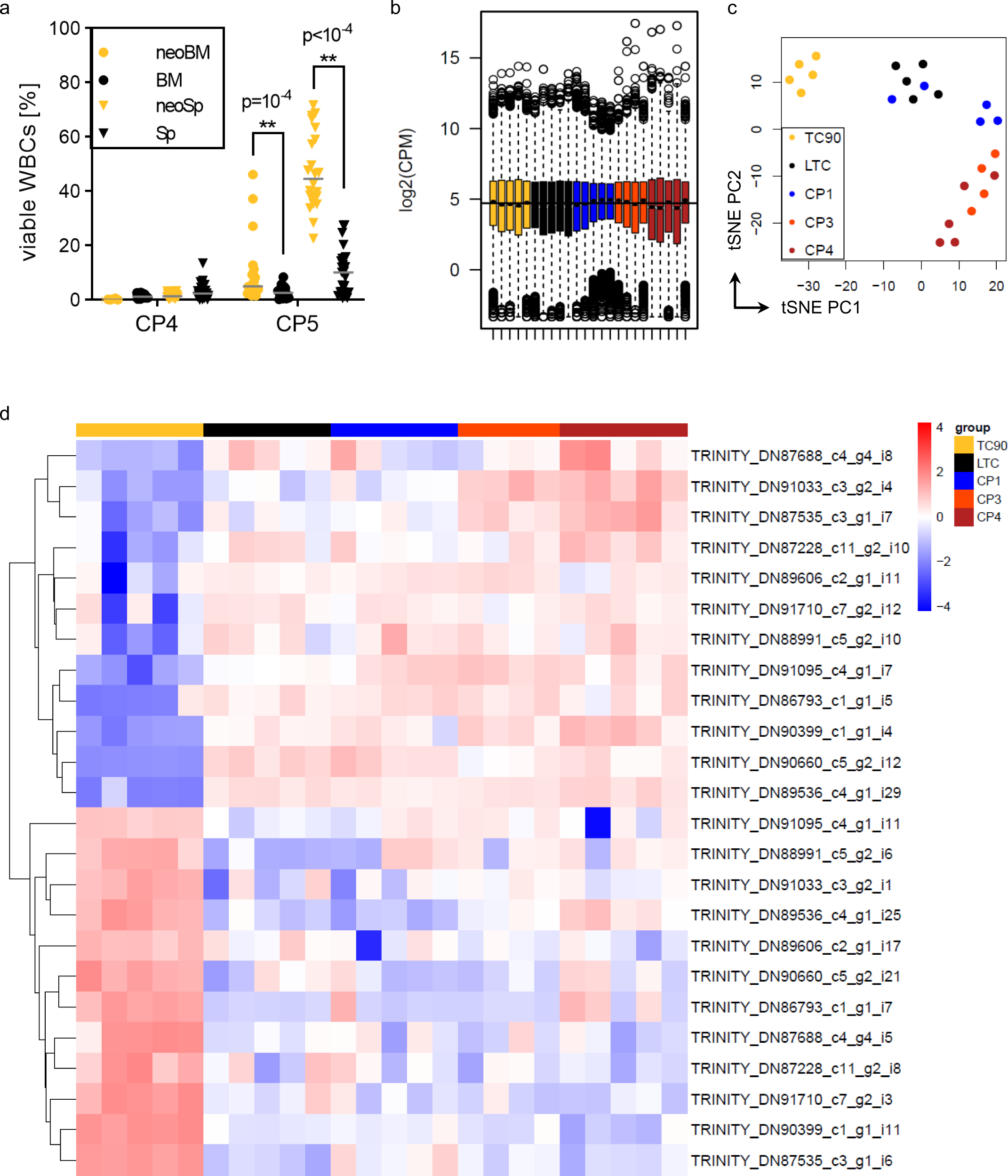
RNA-Seq TC90 vs adult naked mole-rat HSPCs. **a**, Frequencies of CP4 and CP5 across adult BM (N=48), neonate BM (N=23), adult spleen (N=30) and neonatal spleen (N=22); p-value determined by Tukey’s Two-way ANOVA. **b**, Filtered, normalized and log2-transformed naked mole-rat transcriptomes, color code see **c**, Unsupervised t-SNE clustering of neonate TC90 (N=5) and adult LTC (N=5), CP1 (N=5), CP3 (N=4), CP4 (N=5) isoform counts. The clustering pattern of gene counts in Figure 3a is reproduced: LTC/CP1 and CP3/4 form overlapping partitions, TC90s cluster as a distinct group. **d**, Heatmap of 12 inversely expressed transcript pairs from the same gene locus, whereof each single isoform is differentially expressed in TC90 vs other CPs.

**Supplementary Fig. 3.**
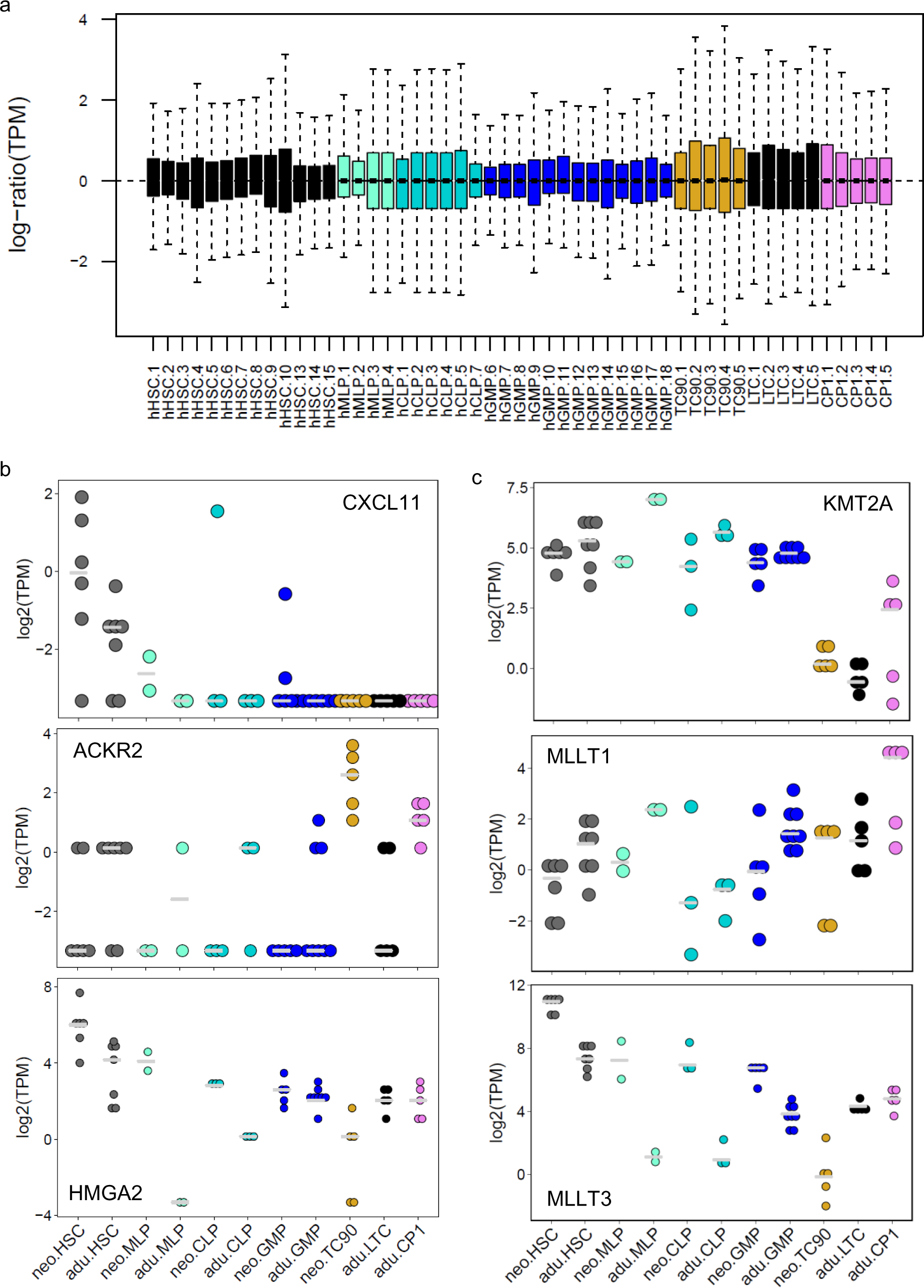
RNA-Seq cross-species, cross-ontogenies. **a**, RLE-plot of filtered, size factor-normalized, log2-transformed and median-centered TPMs [transcripts per million] of human and naked mole-rat transcriptomes. **b**, Log-transformed TPM expression across cell types, species and ontogenies for indicated orthologs.

**Supplementary Fig. 4.**
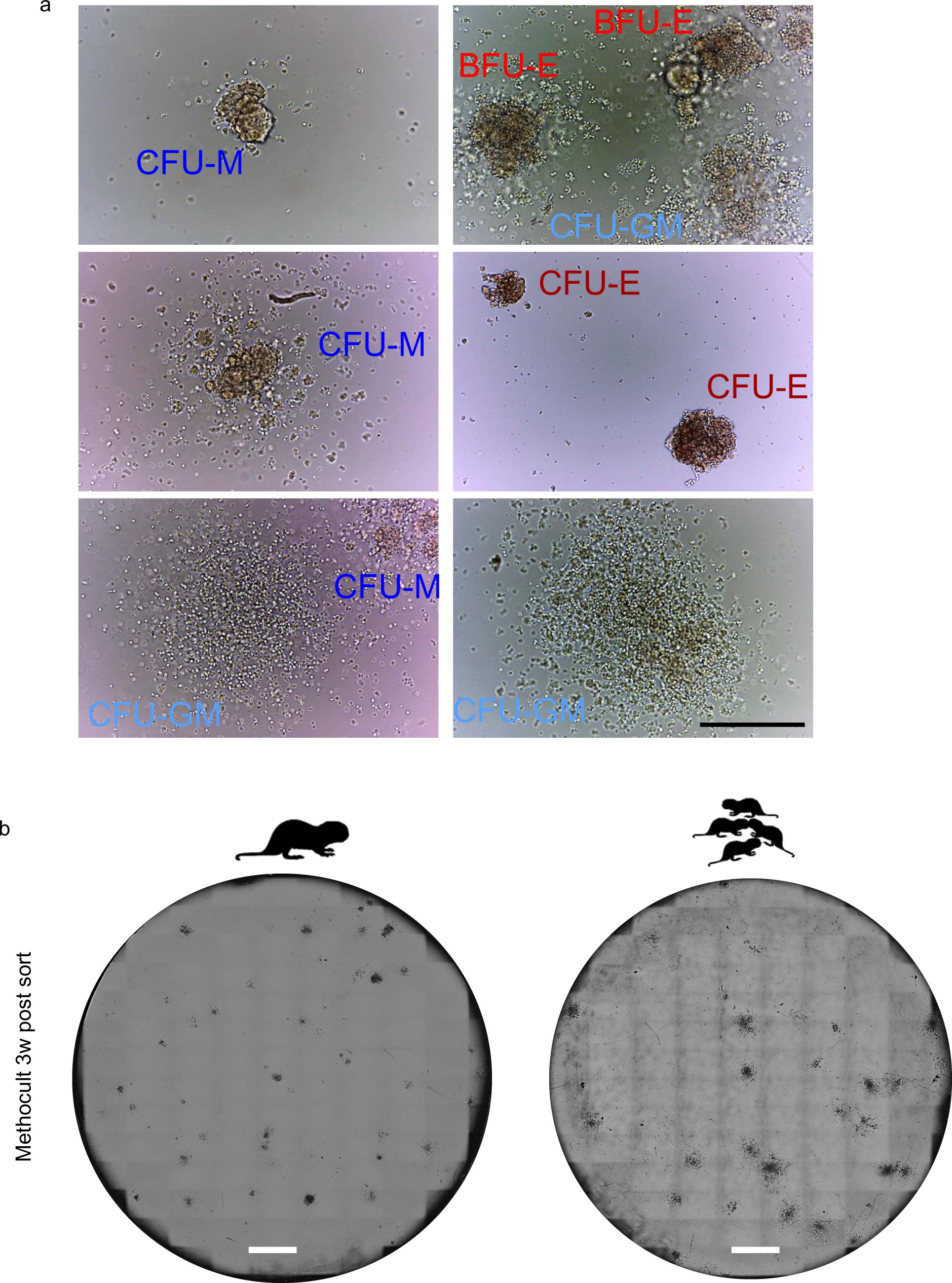
Naked mole-rat colony assays. **a**, Photomicrographs of representative CFU morphologies from an adult BM LTC sample, grown for 3 weeks at 32°C; Scale bar 250µm. Hematoxylin & Eosin staining of spleen sections; Scale bar 200µm [bottom]. **b**, Merged Scan of representative colony assays for adult CP1 [left] and neonate TC90 [right]; Scale bar 400mm. Note that TC90 features larger scattered colonies, pointing towards granulocytic precursors with higher regenerative potential than adult CFU-GMs.

